# Early impairment in dentate pattern separation in a rodent model of temporal lobe epilepsy

**DOI:** 10.1101/2025.11.12.688087

**Authors:** Laura A. Ewell, Ario Ramezani, Gabrielle Suarez, Verónica C. Piatti, Vicky Lam, Tatyana G. Mills, Anelah McGinness, Raafat Kuk, Mika Ukai, Anna-Lena Schlenner, Christopher K. Luong, Stefan Leutgeb, Jill K. Leutgeb

**Author notes:** Correspondence (J.K.L), (S.L.).

## Abstract

The dentate gyrus of the hippocampal formation utilizes sparse neural coding to support memory formation. In temporal lobe epilepsy (TLE), the dentate undergoes extensive restructuring, including loss of local inhibition. It is unknown how circuit changes impact dentate network computations in awake behaving animals. Here we perform high-density tetrode recordings from the dentate in male rats treated with systemic kainic acid (KA) to induce TLE. In KA-treated rats, dentate spatial representations were less precise at the single neuron level, however the stability of the spatial code in representing repeated visits to the same location was retained at the population level. Despite spared spatial representation, the network computations for pattern separation were disrupted. The reorganized dentate network in KA-treated rats generated less distinct population activity patterns for different environments, specifically as a result of impaired rate-remapping. Changes in network computations paralleled impairments in a dentate-dependent memory task, potentially providing a link between altered pattern separation and behavioral discrimination. These deficits were present in animals with confirmed dentate circuit reorganization that had not yet developed spontaneous seizures – suggesting that the dentate network changes underlying altered rate-remapping and impaired dentate-dependent memory occur early in epileptogenesis.

## INTRODUCTION

A critical feature of episodic memory is the ability of neural networks in the hippocampus to represent different events uniquely such that they can be stored and later remembered with minimal interference from memories that share common features. Accurate memory retrieval is thus dependent on encoding memories distinctly when they are formed.

The neuronal mechanism proposed to mediate this process is pattern separation, in which the pattern separator receives overlapping input representing similar experiences, and transforms those inputs to non-overlapping, separated outputs such that a memory is represented by a distinct pattern of activity ^1–3^. Experimental evidence converges to suggest that the first processing station of the hippocampal trisynaptic circuit, the dentate gyrus, is an optimal pattern separator, confirmed both in rodent models ^2,4–7^ and in humans ^5,8,9^. The dentate gyrus is well positioned to perform this operation based on unique network cytoarchitecture, plasticity, and neuronal activity ^10,11^. The dentate gyrus receives dense cortical input from higher cortical areas and sends output (mossy fibers) to the downstream CA3, the auto-associative recurrent network at the core of the hippocampal formation which operates most efficiently for memory encoding when incoming patterns are maximally separated by the dentate gyrus ^1,12^.

At the heart of the capacity for the dentate gyrus neural network to perform pattern separation is its unique sparsity unlike other neural networks in memory circuits of the brain. An extremely sparse neuronal population code is utilized to separate experiences ^13,14,15^. Such sparse activation selection is achieved in part by the large network expansion between the cortex (relatively low numbers of neurons in layer 2 of entorhinal cortex) and the much larger dentate gyrus network ^16^ . Another essential component is strong feedforward ^17^ and feedback inhibition ^18^ that are present within dentate gyrus microcircuits, which effectively silence the majority of neurons, enabling a highly competitive winner take all coding scheme ^19^ and support sub-second toggling between distinct co-active ensembles ^20^.

Further capacity to generate unique population activity is achieved by combining a distributed cell activation code with a distributed rate code for representing distinct experiences. The active subset of dentate neurons vary their firing rates (rate-remapping) or spatial receptive fields (spatial remapping) to represent different features of similar experiences^2^. These collective neuronal changes magnify memory capacity by expanding the possible unique combinations of neuronal activity at the population level ^21,22^. These network dynamics were described in *in vivo* electrophysiological recordings in the dentate gyrus examining the neural mechanism of pattern separation in rodents ^2,4,6^, and can serve as important windows into memory processes. With this mechanistic understanding, the neuronal mechanisms for pattern separation can be assessed as a readout of proper memory encoding and storage of precise spatial memory.

There are several lines of evidence indicating that reduced sparsity in the dentate gyrus weakens pattern separation and leads to memory deficits ^23–26^. Perhaps the most significant line of evidence comes from temporal lobe epilepsy (TLE), a disease defined by increased neuronal activity and synchrony, and thus reduced sparsity. Patients with TLE are unable to pattern separate similar visual scenes ^27^ and mouse models of TLE exhibit reduced *in vitro* dentate pattern separation ^28^. Several changes occur in the dentate gyrus network in epilepsy that could account for dentate-dependent memory deficits. Granule cells sprout de novo axons which create aberrant connections between granule cells ^29–31^. Such increased recurrence has been hypothesized to promote hypersynchronous activation of granule cells, promoting pathological high frequency events ^32^. Other changes include death of inhibitory interneurons ^33,34^, which lead to reduced inhibition onto granule cells ^35^. Finally, granule cells exhibit changes in dendritic and synaptic properties which alter the plasticity rules of the few active neurons ^36,37^. Together, these changes can facilitate overactivation of dentate granule cells ^38^, which leads to runaway excitation and seizure propagation ^39^. It is currently unknown how changes in (1) network recurrence, (2) network inhibition, and (3) seizure burden, impact the neural-network computations in the dentate gyrus that underlie pattern separation and ultimately memory. To address these questions, we first assessed circuit level structural changes and dentate-dependent memory ^40,41^ in rats with ongoing epileptogenesis prior to seizure generation as well as in rats with chronic epilepsy. At the same time-point in disease progression, we performed high-density single unit recordings from the dentate gyrus in a task where the network mechanisms of pattern separation by place field rate-remapping have been extensively characterized. We report that even prior to developing seizures, rats had impaired dentate-dependent memory. This behavioral deficit emerged at a timepoint when we show that the dentate network experienced loss of inhibitory neurons, altered spatial representations and reduced specificity in the population code via reduced rate-remapping between similar environments.

## RESULTS

### Kainic acid induced status epilepticus results in structural changes of the dentate gyrus and impaired behavioral discrimination

To test how circuit reorganization in the dentate gyrus network as a result of epileptogenesis and eventually epilepsy impacts dentate-dependent behavior, we induced *status epilepticus* in male rats utilizing the low-dose systemic kainic acid (KA) model while littermates served as controls. Two months after rats were treated with KA, they were continuously video monitored (24 hours a day) for one month to determine disease progression (those with >2 spontaneous behavioral seizures were classified as epileptic) and then were sacrificed for histological analysis. At this timepoint, roughly half of the treated rats were epileptic, which allowed us to track which changes in the dentate gyrus were associated with seizures (epilepsy) versus associated with epileptogenesis. In both control and KA-induced rat populations, we measured two hallmarks of epilepsy that could have direct impacts on dentate network sparsity; mossy fiber sprouting and hilar somatostatin positive (SST+) interneuron cell loss (Figure 1A-D). Mossy fibers are the excitatory output axons of dentate granule cells and SST+ neurons are interneurons that provide feedback inhibition to granule cells. Mossy fiber sprouting was elevated in KA-treated rats that suffered behavioral seizures, but not in those that were seizure free (Figure 1A,C; Figure S1) (ctrl: n = 28 rats, mean ± S.E.M., 0.7 ± 0.1; KA-no seizures: n = 15 rats, 1.1 ± 0.2; KA-seizures: n = 13 rats, 2.1 ± 0.2; one-way ANOVA, df = 2, f = 18.6, p = 7.5 * 10^-7^, multiple comparisons: ctrl vs KA-no seizures, n.s., ctrl vs KA-seizures, p = 3.9 * 10^-7^, seizures vs no seizures, p = 0.0006). We processed a subset of the tissue to stain for SST+ neurons. The number of SST+ neurons in the dorsal hilus were reduced in both groups of KA-treated rats, regardless of whether they had yet developed spontaneous behavioral seizures (Figure 1 B,D; ctrl: n = 10 rats, mean ± S.E.M., 2525 ± 271 SST+; KA-no seizures: n = 6 rats, 1183 ± 268 SST+; KA-seizures: n = 7 rats, 1371 ± 110 SST+, one-way ANOVA, df = 2, f = 9.52, p = 0.001, multiple comparisons: ctrl vs KA-no seizures, p = 0.005; ctrl vs KA-seizures, p = 0.0009, seizures vs no seizures, n.s.).

**Figure 1.**
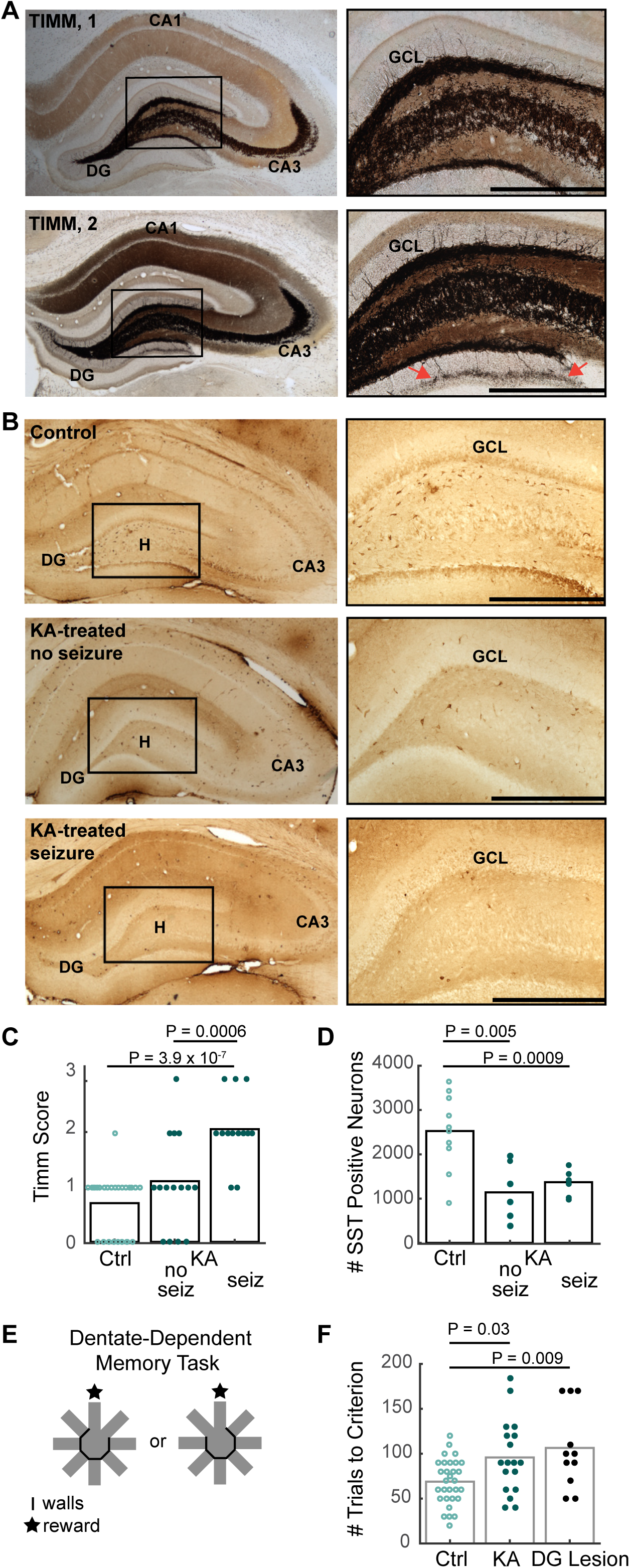
Dentate circuit reorganization and dentate-dependent memory impairment after kainic acid treatment. A, Coronal sections of hippocampi from two KA-treated rats with different degrees of axonal sprouting. Timm stains label zinc concentration black, which is selectively dense in the mossy fiber axons of dentate granule neurons. Magnified images to the right are selected from the area defined by the black boxes centered in the hilar region of the dentate gyrus in the images to the left. Top row, minor sprouting (Timm score 1). Bottom row, intermediate sprouting (Timm score 2). Arrows indicate an example area of sprouting in the inner molecular layer. See Figure S1 for more examples. Scale bars = 250 μm. DG: dentate gyrus; GCL: granule cell layer. **B,** Representative coronal sections of the dentate hilar region from a control rat (top), KA-treated rat without seizures (middle) and a KA-treated rat with seizures (bottom). Somatostatin positive (SST+) neurons are visualized using cell type specific immunohistochemistry. Magnified images to the right are selected from the area defined by the black boxes. (Right) Examples of more abundant SST+ interneurons in the hilus (H) of control rats compared to both KA-treated conditions. **C,** Timm scores indicating degree of mossy fiber sprouting for all rats from each group. Each circle is the average score from one rat. KA-treated rats with seizures have the most extensive mossy fiber sprouting as indicated by significantly higher TIMM scores compared to control rats and KA-treated rats without seizures. **D,** The number of SST+ interneurons in the dorsal hilus is significantly higher in control rats compared to KA-treated rats with and without seizures. Each circle is the average across sections from a single rat. **E**, (Left) Schematic of the dentate-dependent discrimination memory task on an 8-arm radial maze. A single arm is baited with a reward, and two arms are available to the animal, the baited arm and one of the two adjacent arms. The maze was in a small room with distinct distal cues. **F,** The number of trials for rats to preferentially enter the rewarded arm compared to adjacent non-rewarded arms (criterion: 2 of 3 days with ≥90% rewarded trials) by groups (control, KA-treated, and dentate lesioned animals). KA-treated rats with and without seizures are combined, as there were no significant differences between these groups. Ctrl, control; KA, kainic acid treated; seiz, seizures.

A subset of the KA induced animals (n = 18) were tested on a dentate-dependent spatial discrimination task during the period that they were being video monitored. In this reference memory task, on each day of training animals choose between two adjacent open arms of an 8-arm radial arm maze, one of which is consistently baited with a food reward (Figure 1E). To confirm dentate dependence of this task, we added an additional group of animals in which we lesioned the dentate gyrus ^42^. Dorsal dentate granule cells were selectively lesioned with bilateral infusions of colchicine in the dorsal region of the dentate gyrus of the hippocampus. In the lesioned rats (n = 11), there was a 67.3 ± 4.9% reduction in the volume of the dorsal dentate gyrus granule cell layer, as quantified using the Cavalieri method. We found that lesioning the dentate was sufficient to disrupt learning (requiring more trials to reach criterion) and strikingly, KA-treated rats behaved as poorly as the lesion group (Figure 1F) (ctrl: 70 ± 5 trials, n = 30 rats; KA: 95 ± 10 trials, n = 18 rats; DG Lesion: 105 ± 15 trials, n = 11, one-way ANOVA, df = 2,f = 6.27, p = 0.003; multiple comparisons: ctrl vs KA, p = 0.03, ctrl vs lesion, p = 0.009, KA vs lesion, p = 0.7). Within the KA group, there was no behavioral difference between KA animals with or without seizures (seizures: 110 ± 20 trials, n = 7 rats; no seizures: 90 ± 10 trials, n = 11, unpaired t-test, t = 1.19, df = 16, p = 0.25), suggesting that brain changes associated with epileptogenesis were sufficient to disrupt dentate-dependent memory. Collectively, these data imply that early changes to the dentate gyrus network during disease progression, i.e. before the emergence of the spontaneous seizures that define epilepsy, are sufficient to alter the network computations necessary to support behaviors that require fine spatial discrimination.

### Impaired spatial information and reduced spatial sparsity in the neural activity of KA-treated rats

To link how disease related circuit reorganization in the dentate gyrus resulted in altered dentate-dependent memory we next focused on characterizing changes in dentate network activity in KA treated rats. We first asked whether structural changes such as loss of inhibition and increases in excitatory connectivity in epileptogenesis are accompanied by physiological changes in dentate neurons in awake-behaving rats. We recorded activity from dentate principal neurons while rats explored distinct spatial environments, a condition where dentate neurons in the healthy brain were previously shown to represent different environments with unique firing patterns ^2,4,6^. Rats were trained to forage for food in open-field environments that differed in shape (Ctrl: n = 39 neurons, n = 6 rats; KA: n = 60 neurons, n = 6 rats). We tested whether there were general differences in dentate cellular excitability between control animals and those that had undergone KA induced *status epilepticus* 2 months prior.

Distributions of peak firing rates during foraging were similar between the two groups (Median, IQR; Ctrl: 2.2 Hz, 0.3 - 5.6 Hz; KA: 2.0 Hz, 0.5 - 7.8 Hz). Neurons with peak firing rates >2 Hz were considered active during foraging and comprised a similar proportion of total neurons in control and KA-treated rats (Ctrl, n = 22 neurons or 56%; KA, n = 30 neurons or 50%; Figure 2B). Of the active neurons (>2 Hz), KA-treated dentate neurons had similar average firing rates (M, IQR: 0.9 Hz, 0.6 - 1.7 Hz) compared to control neurons (M, IQR: 0.3 Hz, 0.2 - 0.6 Hz; p = 0.13, rank-sum; z-val: -1.5; Figure 2C). Despite similar firing rate distributions, we observed differences in the spatial receptive fields of active cells, measured by assessing the quality of their spatial representations while rats explored the open field. In control rats, even neurons with low spatial information scores (a measure of how well a neuron’s firing rate predicts the rat’s position) exhibited well defined place fields, the difference to higher scores being that cells with lower spatial information had multiple rather than single well defined firing fields within the same environment (Figure 2D, left, bottom row). In contrast, in KA-treated rats, cells had spatial information scores that were below the control range and the cells with the lowest scores were characterized by distributed firing patterns such that the neurons fired across most spatial bins in the open-field (Figure 2D, right, bottom row). In line with this observation, across the population of active neurons, spatial information was lower in KA-treated rats (M, IQR: 0.8, 0.4 - 1.7) compared to control rats (M, IQR: 1.8, 1.2 - 2.2; p = 0.0056; rank-sum, z-val: 2.8; Figure 2E, left). Furthermore, we observed differences in spatial non-selectivity (a measure of the proportion of spatial bins within the environment that the neuron was active in). Across the population, KA-treated dentate neurons exhibited higher spatial non-selectivity (KA-treated neurons: M, IQR, 0.42, 0.19 - 0.60 compared to control neurons: M, IQR, 0.19, 0.15 - 0.29; p = 0.017, rank-sum, z-val: -2.4; Figure 2E, right). These results confirm that the spatial firing patterns of dentate neurons following epileptogenesis are altered despite retained average firing rates. Together, the reduced spatial information and increased spatial non-selectivity indicate that in KA-treated rats, at each location within an open-field, a higher number of dentate neurons are active. Therefore, during an experience at any given moment in time, the dentate network is indeed less sparse in KA-treated rats in comparison to controls. We observed behavioral seizures (Racine III-V) in four of the six KA-treated rats (4 hrs/day of video monitoring, see methods for more details). We therefore asked whether the observed changes in place field quality were driven by neurons recorded in animals with observed behavioral seizures. When testing between neurons recorded from KA-treated rats with seizures (n = 17) vs. without seizures (n = 13), we found no differences in average firing rate (with seizures: 0.6, 0.3 - 1.6; without seizures: 0.3, 0.2-1.2, rank-sum, p = 0.4, z-val:-0.8; Figure S2A), spatial information (with seizures: 0.8, 0.4 – 1.7; without seizures: 0.8, 0.6-1.3, rank-sum, p = 0.77, z-val: 0.29; Figure S2B), or spatial non-selectivity (with seizures: 0.5, 0.2-0.7, without seizures: 0.4,0.3-0.5, rank-sum, p = 0.62, z-val: -0.5; Figure S2C), suggesting changes in neuronal firing patterns occurred prior to seizure generation. We therefore pooled all KA-treated animals for further electrophysiological analysis.

**Figure 2.**
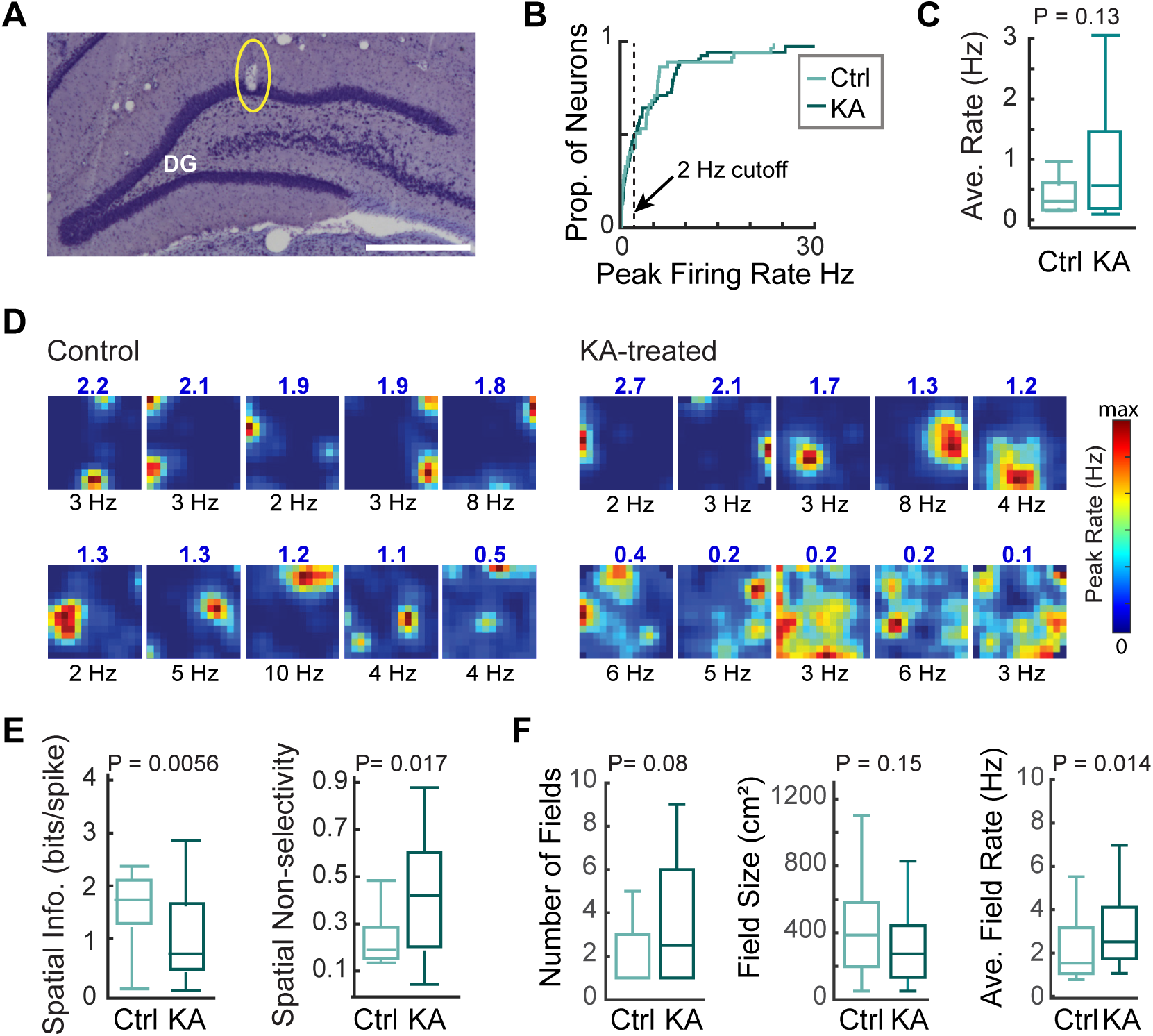
Spatial firing patterns of dentate neurons were less precise in KA-treated rats. A, Cresyl stained coronal section of the dentate gyrus, from a representative control rat with chronic electrophysiological recordings during behavior. The end of a tetrode positioned within the granule cell layer (GCL) is indicated by a circle (see Methods for anatomical tracking of tetrode locations). Neurons are defined as putative granule neurons when the associated tetrode is confirmed by histology to be in the GCL. Scale bar = 250 µm. **B,** Cumulative distribution functions of peak firing rates of recorded dentate neurons (from control and KA-treated rats). Neurons with peak activity >2 Hz (indicated by black dashed line) were considered active. **C,** The average rate of active neurons was similar between KA-treated and control rats. **D,** Activity patterns of individual dentate neurons (10 per group) during 10 min random foraging in a square environment. For each dentate neuron, the spatial information (blue number on top of map) and peak firing rate (black number on bottom of map) were calculated. For each group (control, left; KA-treated, right), the top row comprises rate maps from neurons with the highest spatial information and the bottom row comprises rate maps from neurons with the lowest spatial information. **E,** Firing patterns of dentate neurons carried significantly less spatial information in KA-treated compared to control rats. Rate maps of KA-treated dentate neurons contained significantly higher spatial non-selectivity compared to control rats – revealing that spikes were more spatially dispersed after KA treatment. **F,** Number of place fields of a single dentate cell during one 10-min session in the open field and the average size of each place field did not differ between groups. However, the within field average firing rate was higher in KA-treated compared to control rats. Boxplots show median and quartiles, with whiskers extending to minimum and maximum values. *P* values are reported for rank-sum tests. Ctrl, control; KA, kainic acid treated.

One feature of neurons in the dentate gyrus is that individual place cells often have many distinct spatial firing fields. Multiple place fields per cell endow the sparse network with the capacity to create as many distinct population codes with fewer active cells as a much larger network of active cells can generate from fewer place fields per cell ^2^. We observed that the number of fields per neuron and the size of fields were not statistically different between control and KA-treated rats (Figure 2F). Note, however, that we observed a skewed distribution for number of fields in dentate neurons recorded in KA-treated animals, with a quarter of cells having >5 fields. KA-treated rats had significantly higher within field mean firing rates, unmasking a hyperexcitability phenotype that was occluded when considering overall activity (ctrl: n = 24 fields, 2.3 ± 0.3 Hz; KA: n = 55 fields, 3.1 ± 0.2 Hz, rank-sum, z-val = -2.4, p = 0.014; Figure 2F, right). Collectively we observed several features of altered dentate neuronal activity in KA-treated rats that suggests reduced network sparsity manifests early in disease progression before seizures emerge. These results are in line with our observation that dentate-dependent memory impairments were also present in animals treated with KA, but prior to the emergence of behavioral seizures.

### Dentate network computations that encode and represent events distinctly are impaired in KA-treated rats

An important neural-network computation of the dentate gyrus is to distinguish between similar input patterns, or events, by generating distinct population activity for each input that can then be conveyed to downstream hippocampal subregions that are not optimized for pattern separation ^2,43^. Pattern separation is considered a critical network computation for the encoding of memory that facilitates accurate memory retrieval. To differentiate between environments, dentate neurons either change their spatial firing fields, in-field firing rates, or turn on/off such that the neural representations of similar environments are strongly decorrelated from one another at the population level ^21,44^. To test whether dentate network computations for pattern separation were altered in KA-treated animals, we trained rats to explore an open field arena with flexible walls that could be in either a square or circular configuration (Figure 3A). The arena remained in the same spatial location in the room, with walls placed upon the same section of floor leaving the majority of distal and proximal cues unchanged between trials except for the shape of the enclosure walls. Trials consisted of four ten minute-sessions in which box shape varied between a square or circle randomly. This design allowed us to test pattern separation by comparing dentate population activity in the same location between visits when the same environmental cues were repeated (same shape) and when the environmental cues differed (different shape) (Figure 3B).

**Figure 3.**
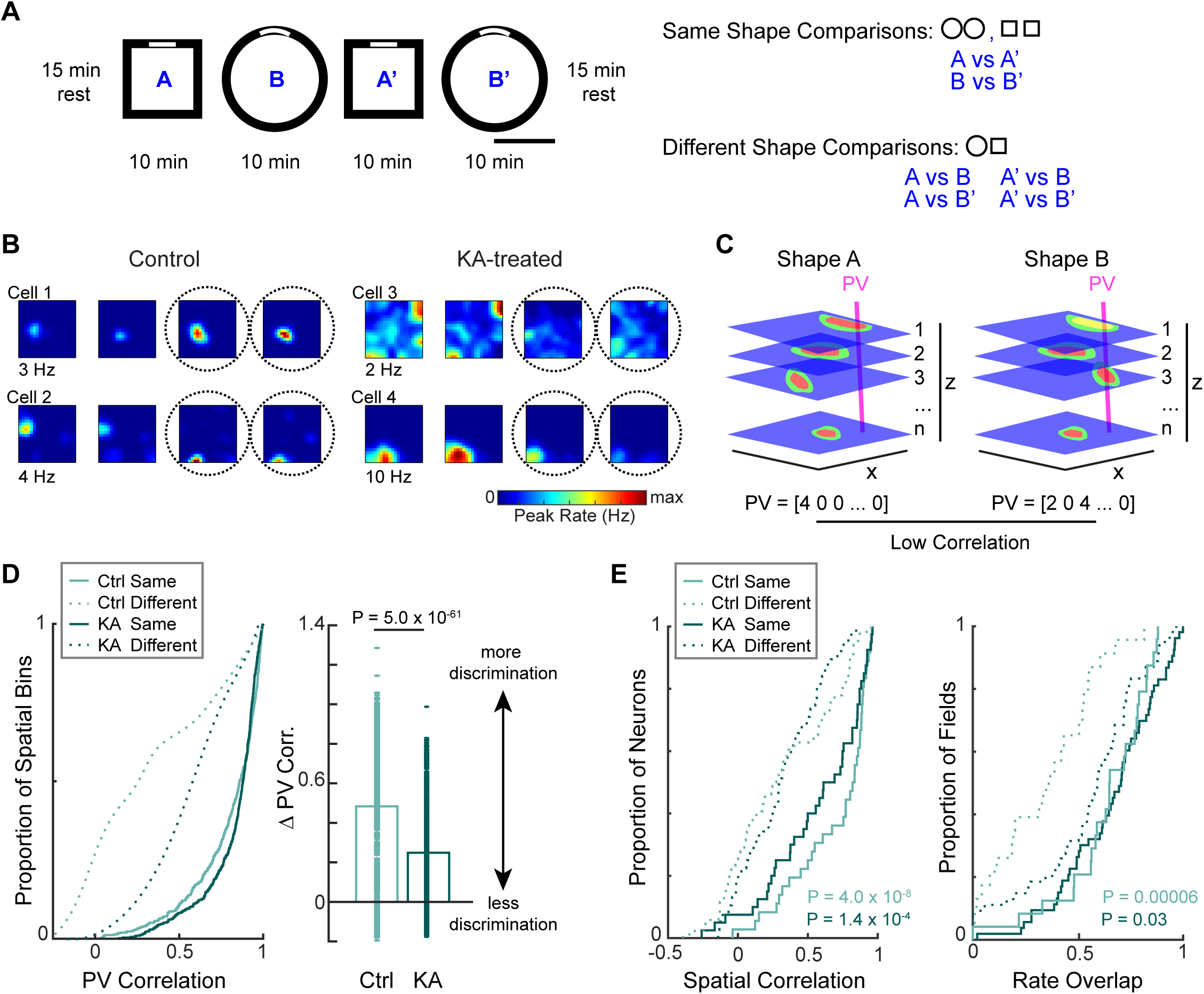
Dentate network pattern separation is impaired in KA-treated rats through reduction in the capacity of the network to perform rate remapping. A, (Left) Behavioral paradigm. Animals explored a highly familiar open-field arena in which the shape of the arena could be changed from square to circular (four 10-min sessions; two of each shape presented in random order each day). The arena remained in the same place in the room with common visual cues and a shared floor. The activity of dentate neuron populations was recorded throughout the entire behavioral paradigm. (Right) Dentate population activity was compared across the behavioral sessions as indicated. **B,** Rate maps for all four sessions are shown for two dentate neurons per group (left, control; right KA-treated). Only common areas between the square and circle shape were compared (corresponding to the color-coded maps, which fill the size of the square arena, but are inside the stippled line for the circular area). Rate maps are reordered for visual comparison, regardless of their order during the experiment. All maps are scaled to the peak rate across the entire recording sequence (indicated below the leftmost rate map for each cell). Representative dentate neurons show similar activity patterns in repetitions of the same shape environment, indicating that spatial representations are stable. Firing patterns across shapes are more different as expected for pattern separation across distinct environments. **C,** Schematic illustrating how population vector correlations are computed from dentate population activity. Using separate stacks for recorded neurons from control and KA-treated rats, population vectors were calculated for each spatial bin (x,y) in the environment across all neurons (z). Example numeric population vectors (PV) corresponding to the bin indicated by the pink line are shown below each stack. Bins that include firing fields that change between environments (rate or location of firing) will have low correlation. **D,** (Left) Cumulative distribution functions of population vector correlations between pairs of spatial bins (comparisons indicated in A, right). The dentate population activity was well correlated between repetitions in same-shaped environments in both control (light colored line) and KA-treated rats (dark colored line). When comparing population activity between different shaped environments (dashed lines), the control group exhibited strong decorrelation (light colored dots), in comparison to the KA-treated group where a more moderate decorrelation was observed (dark colored dots). (Right) Control neuron populations discriminated more between different-shape comparisons versus same-shape comparisons than populations from KA-treated rats (change in PV correlation was calculated by subtracting each different-shape-PV from the median of the same-shape-PV), indicating stronger pattern separation in control networks. **E,** (Left) Cumulative distribution functions of each neuron’s spatial correlations between pairs of firing rate maps (comparisons indicated in A). Same-shape comparisons were more correlated than different-shape comparisons in both groups (KA-treated, control). P values are reported for post-hoc comparison for KA-treated or control between same and different shapes. (Right) Cumulative distribution functions of the rate overlap between pairs of firing fields (comparisons indicated in A). Only the control group demonstrated strong rate-remapping across distinct environments.

To make these comparisons, we constructed population vectors that allow for testing of dentate network decorrelation between environments. Population vectors include both the spatial aspect (where in the environment the neurons are active) and the rate aspect (the magnitude of the activity in each spatial bin) for the entire population of active cells in the network ^21^ (Figure 3C). Place maps from each neuron were generated for each of the four sessions (i.e., two for each of the two environment shapes). The maps of neurons that were active in at least one of the shapes were stacked into a 3-dimensional matrix with x and y corresponding to matching spatial bins and z corresponding to cell identity. Pairs of vectors for each spatial bin(x,y), comprising the average rate for every neuron within that bin, were correlated across either maps from the same shapes (square-square, circle-circle) or different shapes (square-circle). First, we confirmed previous results ^2^ for dentate network computations in control animals (Figure 3D). In control, same shape PV correlations were high, reflecting the same population activity in repeated visits to the same locations (correlation = 0.79 ± 0.009). As expected, different shape comparisons plunged to low correlation values reflecting different patterns of activity associated with different environments (correlation = 0.31 ± 0.01, reduced by 0.48 ± 0.01). In KA-treated rats, same shape PV correlations were also high (correlation = 0.82 ± 0.01), but were reduced to a lesser extent across different shapes (correlation 0.57 ± 0.01, reduced by 0.25 ± 0.01, p = 5 * 10^-61^, unpaired t-test; t stat: 17.05). Thus, the discrimination of different environmental cues was diminished for dentate networks of KA-treated rats compared to control rats.

Next, we further explored the nature of the network discrimination deficits in KA-treated rats to determine whether changes in spatial representations or rate distributions were differentially affected in the KA-treated dentate network. Spatial correlations between the place maps generated from active dentate neurons across same-shape comparisons (circle-circle and square-square) were high for control and KA-treated rats, indicating robust stability of spatial representations for both groups (mean ± S.E.M.; Ctrl: n = 36 comparisons, 0.69 ± 0.04; KA: n = 40 comparisons, 0.57 ± 0.05). Furthermore, a correlation of place cell maps between different environments yielded significantly lower values than for the across-shape comparison for both groups (one-way ANOVA, df = 3, f = 17.39, p = 3.2 * 10^-10^, p<0.001, post-hoc Tukey-Cramer correction for group comparisons, mean ± S.E.M.; Ctrl: n = 76 comparisons, 0.31 ± 0.03, p = 4.0 * 10^-8^; KA: n = 86 comparisons, 0.31 ± 0.04, p = 4.0 * 10^-4^; Figure 3E, left). In contrast to the similar response of spatial representations in control versus KA-treated rats, we observed a difference in rate discrimination by place fields (Figure 3E, right). Note that rate analyses were done on individual place fields rather than the full spatial map of an environment to account for the independent rate code of individual firing fields that have been shown to represent changes in non-spatial cues across environments ^2^. In control rats, the firing rates within place fields were similar between visits to the same environment, but more distinct between visits to different environments, as would be expected when changes in rate is used to discriminate between different environmental cues (reduction of 0.3 ± 0.05, n = 24 fields, sign-rank, z-val = 4.01, p = 5.9 * 10^-5^). KA-treated rats also exhibited a higher rate difference between different compared to same environments (reduction of 0.12 ± 0.04, n = 55 fields, sign-rank, z-val = 2.1, p = 0.03). However, the magnitude of the rate change between same vs. different was significantly larger in controls compared to KA-treated rats (rank-sum, z-val = 3.2, p = 0.0013). Taken together, our results suggest that pattern separation supported by a redistribution of firing rates, a mechanism for the generation of unique population codes for distinct events, is selectively more sensitive to changes in the dentate network as a result of epileptogenesis.

## DISCUSSION

The dentate gyrus undergoes massive restructuring in TLE, yet there have been few studies examining how neural network computations of the dentate gyrus may or may not be compromised. In line with previous work, we observed two histological changes in the dentate gyrus with the induction of epileptogenesis – sprouting of recurrent granule cell axons and loss of somatostatin interneurons – both of which would be predicted to lead to hyperexcitability of dentate gyrus neurons and a loss of network sparsity. In parallel we observed a deficit in dentate-dependent memory along with the degradation of neural-network computations essential for the dentate gyrus contribution to memory encoding. Epileptogenesis induced impairment in the spatial coding features of individual dentate neurons including less precise spatial firing with higher within field firing rates. However, despite a less precise spatial representation at the single neuron level, the stability of the spatial code for repeated visits to the same environment was retained at the population level. The degraded pattern separation in the dentate networks of KA-treated rats for representing distinctly different environments was instead a result of reduced rate-remapping that impaired the capacity of the reorganized dentate network to generate distinct population patterns needed to distinguish between similar familiar environments, which may in turn preclude the cueing of accurate memory retrieval in downstream neural networks.

In our experiments, KA-treated rats showed interneuron loss, behavioral deficits, and impaired pattern separation in the dentate even if they had not yet developed the spontaneous behavioral seizures that define epilepsy. These results suggest that dentate specific memory deficits such as learning to discriminate adjacent spatial locations may be an early manifestation of epileptogenic circuit changes that impair select dentate network computations prior to the development of spontaneous seizures. The timeline in dentate may be different than in other regions of the hippocampus. For example, in CA1, it has been reported that place field spatial information reduced concurrently with the progressive increase in spontaneous seizure frequency and that reductions in spatial stability emerged weeks later ^45^. Earlier impairments in dentate function may provide a biomarker for assessing the risk for developing acquired epilepsy after brain injury or stroke. Indeed, epileptic patients clearly exhibit impaired pattern separation ^27,28^, although to our knowledge no longitudinal study has been done to assess whether performance on such tasks predict disease outcomes.

An early feature of epileptogenesis is interneuron loss in the hippocampus ^46^. Specific to the dentate gyrus, loss of interneurons in the hilus has been widely reported ^35,47^. Even though remaining interneurons sprout, they are unable to compensate, resulting in a net loss of inhibition to the dentate network ^48^. Confirming these reports, we also report a loss of interneurons in the dentate hilus – specifically SST+ containing interneurons (Figure 1D). It is

interesting to consider how this early circuit change may be linked to the altered physiology we observe. Lateral inhibition has been shown to exert a critical circuit level constraint on the number of neurons active at a particular location ^49,50^, which may explain the reduced spatial selectivity we observed in neurons recorded from KA-treated rats. SST+ interneurons are also known to regulate spike bursting within place fields ^51^. Consistent with this role, we observed increased mean firing rate within fields (Figure 2F). The early influence of SST+ cell loss on network rate-remapping could be a combination of both direct effects of the reduced inhibition on select cortical inputs to dentate, as well as the broader effects of reduced inhibition on dentate network sparsity.

Dentate network sparsity is hypothesized to play a critical role in setting the level of pattern separation that can be achieved through unique combinations of population activity ^14^. The fewer neurons capable of overcoming the strong inhibition characteristic of a competitive network like the dentate gyrus ^19^, the more unique patterns of activity can be achieved from the total number of dentate neurons. The possible number of unique population codes is hypothesized to be directly related to the memory capacity of the hippocampus ^1,12^. Moreover, it has been shown experimentally that directly altering the inhibitory gain of the dentate network through the loss of other inhibitory cells results in more neurons becoming active for a given input representation (reduced sparsity) and impaired pattern separation ^52,53^. However, the loss of SST+ interneurons could have a more direct influence on rate-remapping, the identified network computation selectively impaired early in epileptogenesis while spatial coding is retained. Somatostatin interneurons inhibit the distal dendrites of dentate granule neurons in the outer molecular layer ^54^. The outer and inner molecular layers are defined by select cortical inputs that terminate on distinct regions of dendrite. Input from the lateral entorhinal cortex (LEC) reaches granule neurons via the lateral perforant path, terminating on the distal dendrites of granule neurons where SST+ interneurons regulate activity ^55^. Input from the LEC has been shown to convey mainly nonspatial information to hippocampus ^56,57^ and to be necessary for rate-remapping to occur in CA3 neural-networks under the same behavioral conditions tested here ^58^. These results suggest that a select disruption of LEC inputs to dentate gyrus via a reduction in SST+ inhibition could also result in impaired rate-remapping with a preservation of the spatial code from medial inputs. Although direct tests are needed to reveal longitudinal timelines and include causal manipulations, our results suggest a direct link between SST+ loss and an early impairment in dentate network computations for memory rather than spatial coding, providing a therapeutic target for early intervention and the potential rescue of memory comorbidities.

The second histological feature we considered as an early feature of epileptogenesis in the hippocampus was mossy fiber sprouting. We only observed significant levels of sprouting in animals in which we observed spontaneous seizures. The link between sprouting and pathological population bursts in the dentate has been previously reported ^29,59^. Our data suggest that mossy fiber sprouting is better associated with seizure outcomes and is preceded by changes to dentate network computations and dentate-dependent memory deficits. Our findings do not preclude the possibility that sprouting and seizure activity would not further exacerbate cognitive problems. In fact, seizure activity has been shown to impact hippocampal neuronal activity and network computations in additional ways, included changes in spatial information processing ^60^ ^61^ . These results highlight that the disease progression that leads to dentate dependent memory problems versus seizure susceptibility are on separate trajectories and may not share common mechanisms.

Together, these data highlight the importance of the dentate gyrus for pattern separation and underscore its dysfunction in TLE cognitive comorbidities. Impairments in dentate physiology emerge before spontaneous seizures, indicating that functional assessment of dentate memory could be a powerful biomarker for determining disease outcomes. Testing pattern separation in humans has been done at both the imaging and behavioral level ^5,8,9^, further motivating the therapeutic importance for our finding that pattern separation computations in the dentate gyrus are early outcomes of epileptogenesis and precede seizure outcomes. These results also point to a fundamental role of inhibition in the dentate network for supporting its function – indicating that interneuron transplant therapies ^62^, which have had success in treating cognitive comorbidities after brain injury could be an exciting avenue for dentate-dependent memory in TLE as well ^63^.

## Acknowledgements

We thank Y. An, M. Anderson, M. Appalaraju, M. Bangian, D. Esteban, K. Nourmahnad, and M. Wong for technical assistance. This work was supported by the Hellman Foundation, Epilepsy foundation grant 157927, NIMH grant R01 MH119179, and the Walter F. Heiligenberg Professorship to J.K. Leutgeb; NIH grants R01 NS102915, R21 MH100354, R01 NS084324, and R01 NS097772 to S. Leutgeb. L.A. Ewell was supported by NIH training grant F32 MH096526.

## Author Contributions

L.A.E., S.L. and J.K.L. conceived experiments, designed study, and interpreted data. S.L. and J.K.L. managed the project. L.A.E., with assistance from L.M., T.G., and A-L.S, performed electrophysiological experiments. L.A.E., G.S., and A.M. scored seizure monitoring, G.S. and A.M. performed immunohistochemistry and microscopy, V.P. provided dentate lesion data, and A-L.S. quantified the lesion data. L.A.E., G.S., A.M., R.K., and M.K. performed behavioral experiments. L.A.E. analyzed data and prepared figures. L.A.E. prepared the manuscript with feedback and editing from J.K.L. and S.L.

## Ethics Declarations

The authors declare no competing financial interests.

## METHODS

### Subjects

All experimental procedures were performed as approved by the Institutional Animal Care and Use Committee at the University of California, San Diego and according to National Institutes of Health and institutional guidelines. Epilepsy was induced as in previous studies ^64^ ^61^ using a repeated low-dose kainate chronic model of temporal lobe epilepsy ^65^. Male Long Evans rats (n = 52, 40 days of age, Charles River, CA) were treated with kainic acid (KA) (5 mg/kg, i.p., Tocris) each hour until the onset of status epilepticus, which was defined as >10 motor seizures per hour of class IV or V on the Racine scale ^66^. After the seizure induction protocol, the rats were housed individually and maintained on a reverse 12 hr light/12 hr dark cycle. To confirm whether chronic spontaneous seizures that define epilepsy had developed, rats were video-monitored for 4 hours per day for electrophysiological studies or 24 hours per day for behavioral and histological studies. Once two or more motor seizures of class III or greater on the Racine scale were observed, the animal was considered epileptic. Twenty-eight KA-treated rats (3.5 – 4.5 months old) were continuously video monitored for 1 month and paired with 28 control rats not injected with KA (4 months old), after which they were sacrificed for histological analysis of circuit reorganization. Six KA-treated rats (6-12 months old) and six control rats not injected with KA (6 -12 months old) were used for electrophysiological experiments. Two of the control rats used for electrophysiological experiments were wildtype GFAP-TK rats on a Long Evans background ^67^. Eighteen KA-treated rats (6-8 months old) and 30 control rats not injected with KA (6-8 months old) were used for behavioral experiments.

Eleven wildtype GFAP-TK rats on a Long Evans background were injected with colchicine to lesion the dentate gyrus and were used for behavioral experiments. The transgenic GFAP-TK rat line was originally provided by Dr. Heather A. Cameron (National Institutes of Health) and maintained by breeding with wildtype Long Evans rats obtained from Charles River Laboratory.

### Surgical procedures - chronic electrophysiological recording device

Rats were anesthetized with isoflurane (2% to 2.5% in O_2_) and an electrode assembly (Hyperdrive, designed by the McNaughton laboratory ^68^) comprising 14 independently movable tetrodes was implanted above the right hippocampus (AP, 4.1 mm posterior to bregma; ML, 3.0 mm) and fixed to the skull using stainless steel screws and dental cement. Two screws were used as animal ground and were implanted to touch the surface of cortex, anterior and lateral to bregma. Tetrodes were prepared by twisting four insulated platinum wires together (diameter = 0.017 mm, California Fine Wire Company). Leads were plated with platinum prior to surgery to obtain stable impedances near 200 kOhm. Two tetrodes had all four leads shorted with one of the two left in the cortex to record a differential reference signal, while the other was advanced to stratum radiatum to record local field potentials (LFP). The other 12 tetrodes were positioned in the dentate gyrus cell layer to record single units.

### Tetrode recording locations

Histological sections through segments of the hippocampus with electrode tracks were collected as described below (see Histology section). All tetrodes of the 14-tetrode bundle were identified by matching tetrode positions in the bundle with electrode tracks (small lesions) in postmortem tissue. Recordings from a tetrode were included in the data analysis if the tetrode’s deepest position could be clearly identified within or adjacent to (touching) the dentate gyrus granule cell layer (see Figure 2A).

### Surgical procedures - dentate gyrus lesions

Dentate gyrus lesions were performed as described previously ^69^. At the time of surgery, the rat was anesthetized with isoflurane gas [2- 2.5% in O_2_ (20 mL /1 L per minute)] and buprenorphine (0.05 mg/kg, s.c.) was given as an analgesic. To create a lesion specific to granule cells of the dorsal dentate gyrus, colchicine (2.5 mg/ml in 0.1M phosphate buffer, pH 7.2) was injected into 3 sites (0.04 µl per site at 0.012 µl per minute) in both hemispheres at the following coordinates anterior-posterior, medio-lateral from bregma and ventral from dura (in mm): (1) -2.8, ± 0.9, - 3.7 (2) -4.0, ± 2.1, -3.6, (3) -5.2, ± 3.2, -3.7. Three animals received injections only at sites (1) and (2). Infusions were made using a 1 µl Hamilton syringe and the needle was left in place for 8 min at each injection site to prevent spread. Colchicine has previously been shown to cause selective cell death of dentate granule cells ^70^ ^71^. Using computer-based 3D volumetric analysis, the volume of the dorsal dentate gyrus granule cell layer and dorsal CA1 and CA3 pyramidal cell layers was quantified in cresyl violet stained, 40 µm thick brain sections. To assess the completeness of the dorsal dentate gyrus lesions and to assess damage outside the dentate gyrus, the Cavalieri estimator in the Stereo Investigator software (grid size 20 µm; 10x objective) was used for volume quantification. The remaining granule cell volume was normalized to a control volume of 1.54 * 10^9^ µm³ (n = 3 rats). The remaining granule cell percent volume was low and comparable in the two dorsal hemispheres (left: 29.9 ± 4.7 % remaining, right: 35.4 ± 5.7 % remaining). Only minor mechanical damage was observed in the CA1 region above the dentate injection site. Similar damage was previously shown to not significantly contribute to dentate-dependent memory impairment ^69^, or an impairment in CA1-dependent memory ^40^.

### Recordings and data acquisition

Single units and LFP were recorded during restful and running behavior using a data acquisition system (Digital Lynx SX; Neuralynx) with 64 digitally programmable amplifiers. Unit activity was amplified and band-pass filtered between 0.6 and 6 kHz. Spike waveforms above a trigger threshold (40 μV) were time stamped and digitized at 32 kHz for 1 ms. The LFP was recorded continuously in the 0.1 - 900 Hz band from one of the wires of each tetrode. All recording sessions occurred during the animal’s dark cycle (between 7 am - 7 pm). Rats were food deprived to 85% of their baseline weight and trained to forage for randomly scattered food reward (chocolate cereal crumbs) in open arenas. Open arenas were enclosed by a set of flexible walls that could either be in a circular formation (diameter = 1.0 m, height = 0.5 m) or a square formation (0.8 × 0.8 m, height = 0.5 m). Behavioral sessions consisted of four 10 min foraging epochs preceded and followed by resting periods. The rats were also allowed to rest in a holding box for 5 min between foraging epochs. For position tracking, light emitting diodes on the head-mounted preamplifier were tracked at 30 Hz by processing video images.

### Cell clustering

Single units were manually sorted using MClust (MClust 3.5, written by A. David Redish; http://redishlab.neuroscience.umn.edu/MClust/MClust.html) adapted to aid in the tracking of cell identity over extended periods of time ^72^. Clusters that persisted in the same region of parameter space throughout a behavioral session were accepted for analysis, such that the action potentials of a given neuron included in the analysis did not change shape, were distinct from noise signals, and remained inside the cluster boundary. The cluster was required to be clearly defined in both the first and last rest period of the recording session, but was not required to be present in individual foraging epochs, as many dentate gyrus single units are silent within a particular environment.

### Neuronal firing rate map

For each neuron, we constructed firing rate maps for each behavioral session by summing the total number of spikes that occurred in a given location bin (5 × 5 cm), dividing by the total amount of time that the rat occupied the bin, and smoothing with a 5 × 5 bin Gaussian filter with a standard deviation of approximately one bin:

[0.0025 0.0125 0.0200 0.0125 0.0025;

0.0125 0.0625 0.1000 0.0625 0.0125;

0.0200 0.1000 0.1600 0.1000 0.0200;

0.0125 0.0625 0.1000 0.0625 0.0125;

0.0025 0.0125 0.0200 0.0125 0.0025].

Bins that were never within a distance of less than 2.5 cm from the tracked path or with total occupancy of less than 150 ms were regarded as unvisited and were not included in the rate map. Spatial firing patterns of individual cells were compared between sessions when that cell was active (>2 Hz in at least one of the sessions). For each cell, the comparison was performed by calculating the Pearson correlation coefficient for firing rates in corresponding spatial bins across pairs of spatial maps.

### Active neurons

The total number of single units was determined as the number that were tracked across resting and foraging behavior. All included single units were active during the first and last resting period (validating their stability), but not all single units had high firing rates during foraging epochs. Thus, we determine the proportion of active neurons in behavior as the number of neurons that had at least one neuronal firing rate map with a peak >2 Hz divided by the total number of single units recorded (neurons that were clearly isolated and active in sleep sessions before and after running sessions).

### Spatial information score

Spatial information was calculated for each neuron for each foraging session in which the neuron had a peak firing rate of at least 2 Hz in at least one spatial bin. We calculated the spatial information per spike for each firing rate map as: 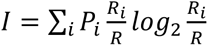 where *i* indexes the spatial bins, P_i_ is the probability of occupancy in each bin, R_i_ is the mean firing rate in each bin, and R is the mean firing rate across the spatial map ^73^.

### Place field boundaries

Individual spatial firing fields of neurons were isolated as previously described ^72^. A reference map was calculated as the average of the common area of the four rate maps corresponding to the four behavioral sessions. For each firing peak in the rate map (>0.8 Hz), contours were calculated at 20 levels between zero and the peak rate. For overlapping contours between fields, each shared contour was divided into segments at inflection points of the contour, and each segment was assigned to the nearest field. After fields were determined, only fields with peak rates >2 Hz and at least 50 mm^2^ were included for further analysis.

### Population vector correlations

For each behavioral session, rate vectors were constructed by arranging the spatial maps of all cells recorded from that region from all animals in an x-y-z stack, where x and y represent the two spatial dimensions and z represents the cell-identity of the population ^44^ ^21^. To allow for comparisons between the square and circular enclosure shapes, the analysis was restricted to the 16 by 16 bins that were common to both shapes, yielding 256 x-y locations. Population vector correlations were obtained by calculating, for each x-y location, the Pearson correlation coefficient for firing rates along the z-dimension between pairs of sessions. Cells with firing below 2 Hz in all x-y bins of the two sessions that were compared were excluded from the population vectors before calculating the correlation coefficients. The correlation coefficients from same shape comparisons (circle-circle and square-square) of all spatial bins were averaged to estimate the population vector correlation for a pairwise comparison between same sessions. The average same shape correlation was subtracted from each different shape comparison correlation to quantify the extent of decorrelation in distinct environments.

### Dentate-dependent behavior

We adopted a behavioral discrimination task that is dependent on the dentate gyrus. Rats learned to discriminate between adjacent arms on an 8-Arm Radial Maze ^42^. The maze was in a room with spatial cues on the walls (not on the maze). The maze was comprised of 8 arms arranged around an octagonal stem. Each arm was spaced 45 degrees apart from its adjacent arms. The maze measured approximately 188 centimeters in diameter and was elevated approximately 36 centimeters above the ground. Transparent Plexiglas walls were placed around the central stem of the maze and attached with Velcro in order to block unused arms and designate the two arms that were open per trial. For the adjacent arm discrimination task open arms would either be arm 1 and 2 or arm 1 and 8. Uniform, chrome, silver bowls were taped to the end of each arm. The bowl of the reward arm (arm 1) was the location of the food reward (chocolate cereal) for all behavior testing performed during this experiment. Rats were food restricted to 85% of their baseline weight and maintained at that weight for the duration of habituation and testing. Habituation was conducted in three phases. The first phase involved placement of a single food reward at the stem of the maze, midway down the three experimental arms, just outside the silver bowls, and inside each bowl. The animal was given 10 minutes to explore during this phase of habituation. If all 10 rewards were eaten within the 10-minute time frame, the rat moved to phase 2. For phase 2, reward was placed within the bowls and just outside the bowls of the same three arms. Eating all rewards within 10 minutes allowed the rat to pass to phase 3, in which reward was only in the three bowls. Rats had 5 minutes to eat all rewards to move onto testing. During testing, the rat ran 10 trials per day. For each trial two adjacent arms were open (1 and 2) or (1 and 8). Each adjacent, non-rewarded arm was used 5 times among the 10 trials in a pseudo-random order to ensure that the rat was not relying on a right-left strategy when making its decision. Reward was always at arm 1. For each day, the number of correct trials were counted. In between trials, the rat was enclosed under an opaque bucket in the central stem. All arms were washed with water and dried between trails. Testing continued until the rat reached a criterion of 90% correct trials for 2 of 3 consecutive days, or for a maximum of 17 days. The total number of trials to reach criterion were summed. Initially, behavioral testing was done in rats that were not monitored 24 hours for behavioral seizures. The behavioral performance of non-monitored KA-treated animals did not differ from video-monitored KA-treated animals (non-monitored: 90 ± 10 trials, n = 8 rats; monitored: 95 ± 10 trials, n = 18 rats; unpaired t-test, t = -0.45, df = 24, p = 0.66), therefore data were pooled. Among monitored KA-treated animals, those that had seizures had similar performance to those who did not (seizures: 110 ± 20 trials, n = 7 rats; no seizures: 90 ± 10 trials, n = 11; unpaired t-test, t = 1.19, df = 16, p = 0.25), therefore data were pooled. Control animals from lesion experiments were 5 months old, while control animals from epilepsy experiments were 6 months old. No statistical difference in behavioral performance was observed between control groups (epilepsy control: 70 ± 5 trials, n = 21 rats; lesion control: 60 ± 5 trials, n = 9 rats; unpaired t-test, t = 1.18, df = 28, p = 0.25), therefore data were pooled.

### Histology

The rat was deeply anesthetized with isoflurane and given a lethal injection of Pentobarbital (0.8 mL - 2 mL). Rats were systemically perfused, first with 0.37% sulphide solution and then with 4% paraformaldehyde (PFA). Brains were transferred to 30% sucrose in 1X PBS before sectioning with a frozen microtome (40 µM sections, Leica).

*Somatostatin immunohistochemistry:* Every sixth, coronal section (40 μm) was processed for staining. Sections were washed (3 x 5 min) in 1X PBS. The free-floating sections were then blocked for 1 hour in blocking solution at room temperature (10% horse serum (HS) and 0.3% Triton-X in PBS). After blocking, the sections were incubated in diluted 1:300 monoclonal mouse anti-somatostatin antibody (GeneTex, code GTX71935, clone SOM-018) with 10% HS and 0.3% Triton-X in PBS over two nights at 4°C. After washing (3 x 5 min) in 1 x PBS, the sections were then incubated for 1 hour at room temperature in diluted (1:200) biotinylated horse anti-mouse IgG antibody (Vector BA-2000) with 0.3% Triton-X in PBS, and then washed again. Endogenous peroxide activity was blocked with 0.3% *H_2_O_2_* for 30 minutes at room temperature. The sections were washed again and then incubated with the avidin-biotin-peroxidase technique (Vectastain ABC Kit, Vector Labs, USA) for 30 minutes at room temperature. The complex was developed using 0.05% of 3,3’-diaminobenxidine (DAB) tetrahydrochloride, 0.3% *NiCl_2_* ꞏ 6*H_2_O* stock, and 0.05% hydrogen peroxide as a substrate for the peroxidase reaction. The sections were then mounted onto Fisherbrand Superfrost Plus Microscope Slides. Stereology of the SST-immunostained tissue was performed using a Leica CTR 6000 microscope and the optical fractionator method and the Cavalieri principle on Stereo Investigator 10 (MBF Bioscience). The counting frame was 300 μm × 300 μm and the counting grid was also 300 μm × 300 μm. After counting SST interneurons in the dorsal hilus of each rat, the total SST interneuron population was estimated based on the thickness of the mounted sections (40 μm).

*TIMM staining:* Sections were mounted onto Fisherbrand Superfrost Plus Microscope Slides with a gelatin and PBS mounting solution and left to dry overnight. Slides were placed in distilled water for 3 minutes and then incubated in developer solution in the dark. The developer solution is: 120 mL of Gum Arabic (100 g/200 mL), 10 mL of filtered citrate buffer (51g *C_6_H_8_O_7_*/200 mL; 47g *C_6_H_5_Na_3_*-*O_7_* - 2*H_2_O*/200 mL), 60 mL of 1.7% hydroquinone solution, and 1 mL of 0.09% light-sensitive, silver nitrate solution. After 30 minutes of incubation, the developer solution was mixed and the tissue was re-submerged for at least another 30 minutes. If the stain was not dark enough, the tissue was left in solution until the stain was complete.

The slides were then dipped twice in warm distilled water (50°C) and placed in room temperature distilled water for 15 to 30 minutes. Next, the slides were fixed in 5% sodium thiosulfat pentahydrat for 30 seconds. Slides were dehydrated in an alcohol series and coverslipped with permount. The tissue for a subset of animals used for electrophysiological experiments were processed for TIMM staining and subsequently stained with cresyl violet to facilitate tetrode tracking ^74^. For TIMM scoring, the scorer was blinded to condition and scored the extent of mossy fiber sprouting according to the criteria described in ^75^. Every sixth section through the dorsal dentate gyrus was given a TIMM score. Sections with no or only occasional supragranular mossy fiber-like staining were scored 0. Sections with scattered mossy fiber-like staining above all parts of the granular layer were given a score of 1. Sections that exhibited either patches of heavy mossy fiber-like staining interspersed with regions of sparser staining or a continuous band of staining intermediate in intensity between sections scored 1 and 3 were given a score of 2. Finally, sections with a dense, continuous band of supragranular mossy fiber-like staining were assigned a score of 3. For analysis, the worst score was selected to represent the maximum extent of sprouting that was observed in a given rat.

### Statistics

For each statistical comparison, normality was assessed with Lilliefors test and either parametric or non-parametric tests were chosen accordingly. Multiple comparisons were made with Tukey Kramer post-hoc tests. Throughout the text values are presented as mean ± S.E.M. or median and inter-quartile interval.

---

**Figure S1.**
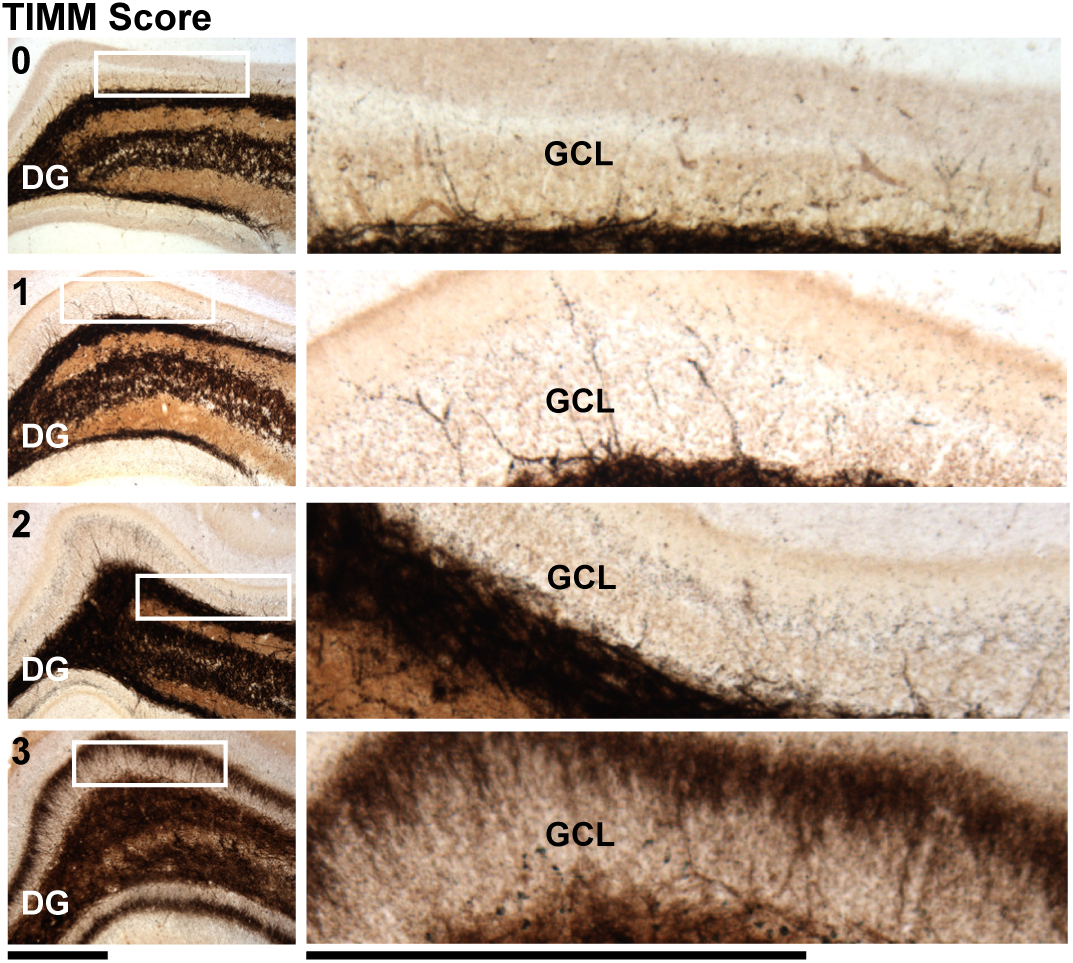
Range of TIMM scores observed in KA-treated rats. (Left) Four images from four different rats, each of which were scored with progressively higher TIMM scores (0-3, top left corner), a measure of the degree of mossy fiber sprouting (see Methods for details on scoring procedure). Sections with no or only occasional supragranular mossy fiber staining were scored 0. Sections with scattered mossy fiber staining above all parts of the granular layer were given a score of 1. Sections that exhibited either patches of heavy mossy fiber staining interspersed with regions of sparser staining or a continuous band of staining intermediate in intensity were given a score of 2. Finally, sections with a dense, continuous band of supragranular mossy fiber staining were assigned a score of 3. (Right) Magnified images from the area defined by white boxes in the sections to the left. The magnified area spans the granule cell layer and inner molecular layer (IML). A higher TIMM score reflects a more substantial reorganization of the mossy fibers, including sprouting into regions where axons and terminals are typically not found, such as the IML. For example, the bottom image with the highest TIMM score of 3 is characterized by extensive black fiber tracks within the IML in comparison to control levels (score of 0). GCL, granule cell layer.

**Figure S2.**
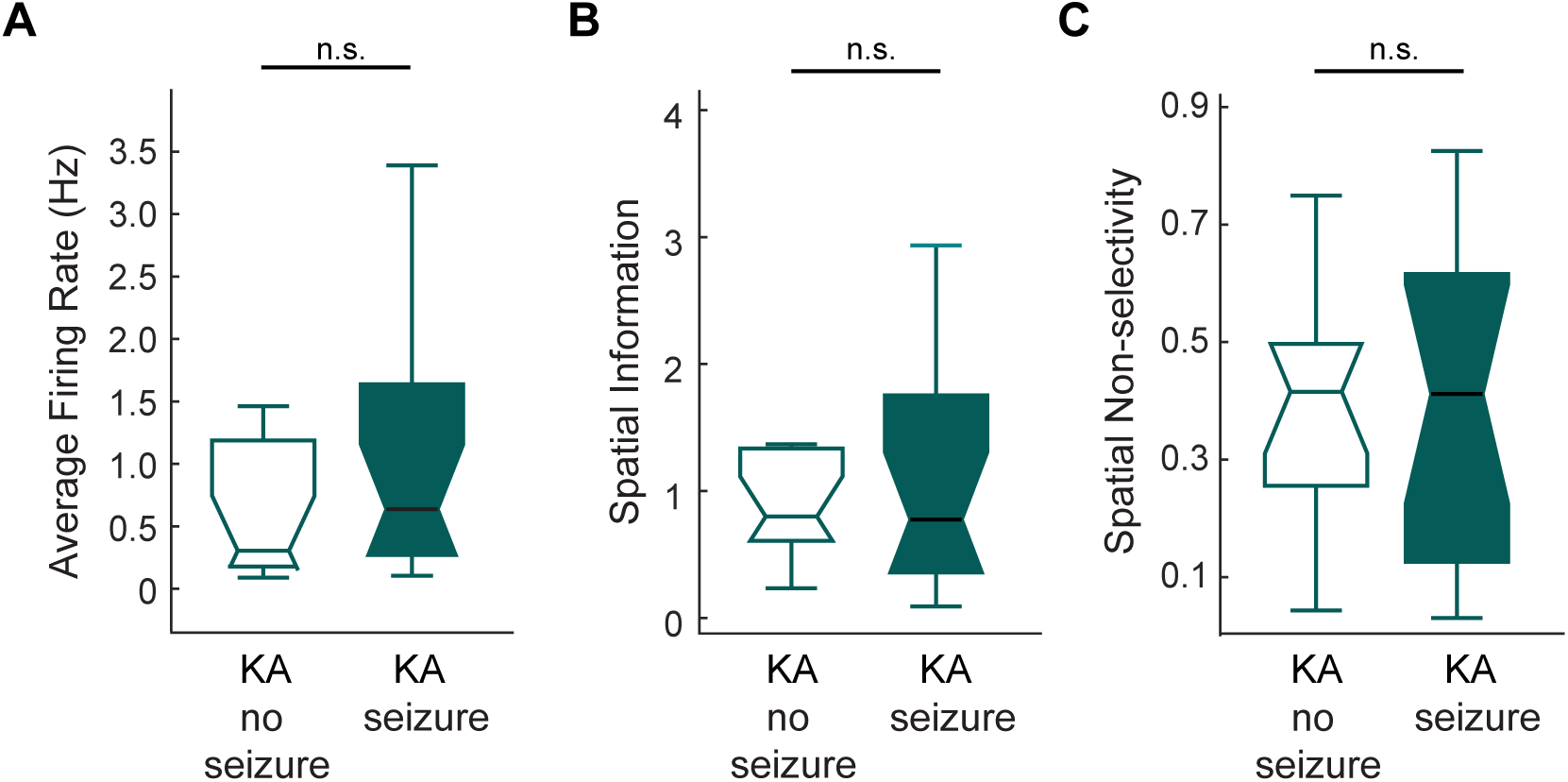
The spatial firing properties of dentate neurons from KA-treated rats without seizures did not differ from rats that had developed epileptic seizures. No statistical differences were measured in (A) average firing rate, p = 0.4, (B) spatial information, p = 0.77, or (C) spatial non-selectivity, p = 0.62, when comparing neurons from KA-treated rats with or without observed seizure events (KA-treated no seizure: n = 2 rats, n = 13 active dentate neurons; KA-treated seizure: n = 4 rats, n = 17 active dentate neurons). Active neurons are defined as neurons with a peak rate >2 Hz. Statistics: rank-sum test. KA, kainic acid treated.

## Notes

### Competing Interest Statement

The authors have declared no competing interest.

### Summary of Updates

This version of the manuscript has been revised to make changes to the author list.

